# The Lon protease temporally restricts polar cell differentiation events during the *Caulobacter* cell cycle

**DOI:** 10.1101/2021.06.29.450348

**Authors:** Deike J. Omnus, Matthias J. Fink, Klaudia Szwedo, Kristina Jonas

## Abstract

The highly conserved protease Lon has important regulatory and protein quality control functions in cells from the three domains of life. Despite many years of research on Lon, only few specific protein substrates are known in most organisms. Here, we used a quantitative proteomics approach to identify novel substrates of Lon in the dimorphic bacterium *Caulobacter crescentus.* We focused our study on proteins involved in polar cell differentiation and investigated the developmental regulator StaR and the flagella hook length regulator FliK as specific Lon substrates in detail. We show that Lon recognizes these proteins at their C-termini, and that Lon-dependent degradation ensures their temporally restricted accumulation in the cell cycle phase when their function is needed. Disruption of this precise temporal regulation of StaR and FliK levels in a *Δlon* mutant contributes to defects in stalk biogenesis and motility, respectively, revealing a critical role of Lon in coordinating developmental processes with cell cycle progression. Our work underscores the importance of Lon in the regulation of complex temporally controlled processes by adjusting the concentrations of critical regulatory proteins. Furthermore, this study includes the first characterization of FliK in *C. crescentus* and uncovers a dual role of the C-terminal amino acids of FliK in protein function and degradation.

## Introduction

Intracellular proteolysis is a critical process in all cell types that is carried out by dedicated proteases. By removing damaged and non-functional proteins, proteases are necessary for maintaining protein homeostasis, in particular under stress conditions that threaten the proteome. Additionally, proteases have important regulatory roles in precisely adjusting the amounts of specific functional proteins, thus complementing transcriptional and post-transcriptional control mechanisms. Because of their important cellular functions, human proteases are considered as promising therapeutic targets [1] and their bacterial counterparts as potential antimicrobial drug targets [2]. Hence, extending the knowledge of the substrate pools of specific proteases and the mechanisms underlying substrate selection is vital.

In prokaryotes and in the mitochondria and chloroplasts of eukaryotes, the majority of proteins is degraded by ATP-dependent proteases of the AAA+ (ATPases associated with various cellular activities) protein family [3]. The protease Lon was the first ATP-dependent protease to be identified and is widely conserved across the three domains of life [4]. Lon forms a hexameric protease complex, of which each monomer contains three functional domains: an N-terminal domain, an ATP-dependent unfoldase domain, and a peptidase domain forming the proteolytic chamber [4]. As a heat shock protein [5], Lon is upregulated in response to protein unfolding stress, such as thermal stress, and contributes to the degradation of un-and misfolded proteins that accumulate under these conditions [4, 6, 7]. In addition to its role in protein quality control, Lon exerts important regulatory functions that can be traced to the degradation of specific sets of native substrate proteins involved in stress responses, metabolism, pathogenicity and cell cycle progression [8].

Despite many years of research on Lon proteases, the number of validated Lon substrates is small in most organisms. Most knowledge about Lon has been obtained by studying bacterial Lon orthologs. In *Escherichia coli* Lon specifically degrades several stress-induced regulators [9–11], metabolic enzymes [12] as well as antitoxins of toxin-antitoxin systems [13], and in several pathogenic bacteria, Lon was shown to degrade regulators of pathogenicity [14, 15], thus playing important roles in the regulation of virulence pathways. In the alpha-proteobacterium *Caulobacter crescentus*, the number of identified Lon substrates to date is small, however Lon is known to impact cell cycle progression by degrading essential cell cycle regulators.

The *C. crescentus* cell cycle is characterized by an asymmetric cell division event and morphologically distinct cell cycle phases [16]. Each division yields a flagellated and piliated swarmer cell and a sessile stalked cell. While the daughter stalked cell initiates DNA replication immediately after cell division, the daughter swarmer cell must differentiate into a stalked cell before entering S-phase. Faithful progression through the *C. crescentus* cell cycle relies on precise coordination of the polar differentiation events that trigger flagella, pili and stalk biosynthesis with core cell cycle events, such as DNA replication and cell division [16]. Previous work established that around one third of all genes in *C. crescentus* show cell cycle-dependent fluctuations in their expression levels [17, 18]. Many of the corresponding proteins have important developmental functions and peak in abundance in the cell cycle phase in which their function is most needed [17]. In addition to transcriptional regulatory mechanisms, active proteolysis must occur to rapidly adjust protein concentrations to enforce these transcriptional changes [19]. However, only a relatively small subset of cell cycle-regulated factors with developmental functions has so far been found to be subject to proteolysis in *C. crescentus* and the contributions of distinct proteases in this process remain incompletely defined.

Previous work established that the ClpP protease with its unfoldases ClpA and ClpX has key roles in *C. crescentus* development by mediating the temporally and spatially controlled degradation of several important cell cycle regulators [20, 21]. In addition to ClpXP, Lon plays an important role in *C. crescentus* cell cycle regulation [20, 21]. It degrades the swarmer cell specific transcription factor SciP [22], the methyltransferase and transcriptional regulator CcrM [23] and the conserved replication initiator DnaA [24]. Lon-dependent degradation of SciP and CcrM contributes to the cell cycle-dependent activity restriction of these regulators [22, 23], while DnaA degradation by Lon ensures rapid clearance of the protein at the onset of nutritional and proteotoxic stress, preventing cell cycle progression under these conditions [24–26]. Although *C. crescentus* cells lacking *lon* are viable, they grow more slowly and show aberrant chromosome content and division defects [23, 25], which can in part be attributed to the stabilization of DnaA, SciP and CcrM. Interestingly, *Δlon* cells exhibit also characteristic developmental defects, i.e., elongated stalks and motility defects [23, 27], suggesting that Lon degrades additional substrates involved in cell differentiation.

Here, using a quantitative proteomics approach, we identified several proteins involved in the dimorphic life cycle of *C. crescentus* as novel Lon substrates. We studied in detail the transcriptional regulator of stalk biogenesis, StaR, and the flagella hook length regulator FliK as specific Lon substrates. We show that Lon is required to establish cell cycle-dependent fluctuations of these regulatory proteins, thereby contributing to their precise temporal accumulation during the cell cycle phase in which their function is needed. Furthermore, we demonstrate that increased abundance of these proteins results in aberrant stalk length and motility defects. Together, our work revealed a critical role of Lon in coordinating cell differentiation with core cell cycle events in *C. crescentus*.

## Results

### A quantitative proteomics approach identifies novel putative Lon substrates involved in ***Caulobacter* development**

Previous work established that Lon degrades the cell cycle regulators DnaA, CcrM and SciP in *C. crescentus* [22–24]. Absence of Lon, either in a *Δlon* deletion strain or following Lon depletion, results in increased stability and abundance of these proteins (Fig. 1A-B). Conversely, *lon* overexpression leads to lower protein abundance of DnaA, CcrM and SciP (Fig. 1C). Based on these results, we reasoned that it should be possible to identify novel Lon substrates by monitoring proteome-wide differences in protein stability and protein abundance in strains lacking or overexpressing *lon* in comparison to wild type cells. Thus, we sampled cells from the following strain backgrounds and conditions for quantitative proteomics analysis: 1) 0, 15 and 30 min following protein synthesis shut-down in the wild type to assess protein stability in the presence of Lon 2) 0, 15 and 30 min following protein synthesis shut-down in *Δlon* cells to examine protein stability in the absence of Lon 3) before and after 4.5 hours of Lon depletion in a *P_van_*-dependent Lon depletion strain and 4) before and after 1 hour of inducing *lon* overexpression from a medium-copy plasmid. Tandem mass tag (TMT) labeling and mass spectrometrical analysis (MS) was used to detect proteome-wide differences in protein levels across these samples. We obtained signals for 2270 or 2261 proteins, respectively, in our two biological replicates and sorted the proteins with respect to four criteria (see Materials and Methods for details): 1) to be more stable in *Δlon* cells than in the wild type after 30 min of translation inhibition 2) to be present in higher abundance at t=0 in *Δlon* cells compared to the wild type 3) to be upregulated after Lon depletion compared to non-depleting conditions and 4) to be downregulated following xylose-induced Lon overexpression compared to non-inducing conditions (Fig. 1D). We identified 26 proteins that fulfilled all four criteria, one of them being DnaA. 120 proteins fulfilled three criteria, and because CcrM and SciP were in this group of proteins (Fig. 1D, Dataset S1), we considered proteins satisfying either three or four criteria as putative Lon substrates. Sorting these 146 proteins by functional category showed that many of them are annotated to have functions in cell cycle and cell differentiation processes (14.4 %), signal transduction (12.3 %), stress responses (14.4 %) and core genetic information processing (6.8 %) (Fig. 1E). Additionally, a large fraction of proteins (17.1 %) is classified as metabolic proteins. Notably, we did not detect FixT and HipB2 in our proteomics experiment (Dataset S1), two other recently reported Lon substrates in *C. crescentus* [28, 29]. Since *Δlon* cells have previously been shown to exhibit developmental defects [23, 27], we focused this study on the group of potential Lon substrates annotated to have functions in cell cycle and cell differentiation processes. This group included proteins involved in central cell cycle regulation, chromosome partitioning, stalk morphogenesis as well as motility and chemotaxis (Fig. 1F). Interestingly, according to previously published RNA-sequencing data, a large subset of the genes encoding these proteins are subject to cell cycle-dependent regulation and peak in their expression during a specific cell cycle phase (Fig. 1F) [30, 31].

**Figure 1.**
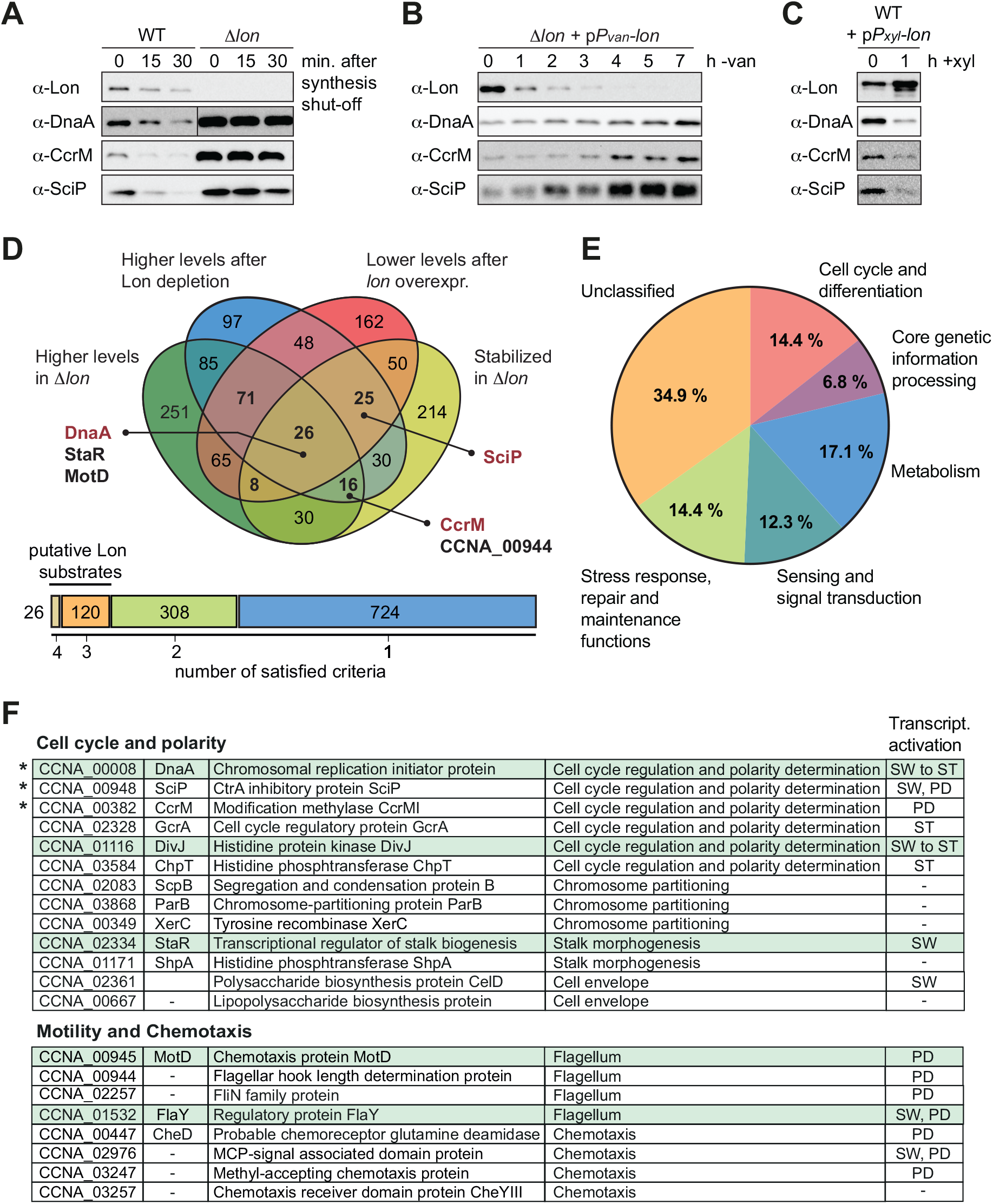
A quantitative proteomics approach identifies novel Lon substrates in *C. crescentus*. (A) *In vivo* stability assays of Lon, DnaA, CcrM and SciP in the wild type and the *Δlon* strain (LS2382). Protein synthesis was shut down by addition of chloramphenicol at t = 0 and remaining protein levels were measured after 15 and 30 minutes. The protein levels at t = 0 correspond to the steady state levels of these proteins in the wild type and the *Δlon* (LS2382) mutant. (B) Protein levels of Lon, DnaA, CcrM and SciP over seven hours of Lon depletion. The expression of *lon* was shut off by transferring the *P_van_*-dependent Lon depletion strain (ML2022) from PYE supplemented with vanillate (van) to PYE lacking vanillate (-van). (C) Protein levels of Lon, DnaA, CcrM and SciP in a *P_xyl_*-dependent *lon* overexpression strain (ML2010) before (0) and 1 hour after induction with xylose (1 h + xyl). (D) Venn chart showing groups of proteins that meet the following criteria and how they overlap: 1) to be present in higher abundance at steady state in *Δlon* cells compared to the wild type (green circle), 2) to be upregulated after 4.5 h of Lon depletion compared to non-depleting conditions (blue circle), 3) to be downregulated after 1 h of induced *lon* overexpression compared to non-inducing conditions (red circle) and 4) to be stabilized in *Δlon* cells compared to the wild type 30 min after translation shut-off (yellow circle). Previously confirmed Lon substrates (shown in red) and the proteins investigated in this study (shown in black) are highlighted. The bar graph below the Venn chart indicates the number of proteins satisfying four, three, two or one criteria. Proteins satisfying either three or four criteria were considered as putative Lon substrates. (E) Putative Lon substrates identified in (D) sorted by functional category. (F) Tables listing the putative Lon substrates with functions in cell cycle and polarity as well as motility and chemotaxis. Previously known Lon substrates are marked with an asterisk. Proteins satisfying four criteria are highlighted in green. The cell cycle phase in which the expression of the listed proteins is transcriptionally induced is indicated (SW: swarmer cell, ST: stalked cell, PD: predivisional cell, SW to ST: swarmer to stalked cell transition).

### The developmental regulator StaR is a specific Lon substrate

One of the proteins that satisfied the four criteria in our proteomics experiment most clearly was the transcriptional regulator StaR, a protein previously reported to be involved in the regulation of stalk biogenesis and holdfast production (Fig. 1D, F) [32, 33]. Like the three known substrates DnaA, CcrM and SciP, StaR was more stable in the *Δlon* strain compared to the wild type and showed increased steady-state levels in *Δlon* and Lon-depleted cells but reduced protein levels in cells overexpressing *lon* (Fig. 2A, Dataset S1). To validate these proteomics data, we monitored the stability of StaR in wild type and *Δlon* cells using a StaR-specific antibody [33]. In the wild type, the levels of StaR were below the limit of detection, however, we observed a strong and stable band corresponding to StaR in the *Δlon* strain (Fig. 2B), indicating that StaR is upregulated in *Δlon* cells, a result that is consistent with the proteomics data (Fig. 2A, Dataset S1). To analyze the rate of StaR degradation in the presence of Lon, we expressed *staR* from a medium-copy vector to elevate its levels. In this strain, StaR was degraded with a half-life of approximately 12 minutes, when Lon was present (Fig. 2C). Absence of Lon resulted again in complete stabilization of StaR and increased steady state levels, which is in line with our proteomics data, and confirms that StaR degradation depends on Lon. To directly test if StaR is a Lon substrate, we performed an *in vitro* degradation assay with purified StaR and Lon. This assay showed that Lon readily degrades StaR in an ATP-dependent manner (Fig. 2D). Hence, StaR is a Lon substrate and no additional factors are required for recognizing and degrading StaR, at least *in vitro*.

**Figure 2.**
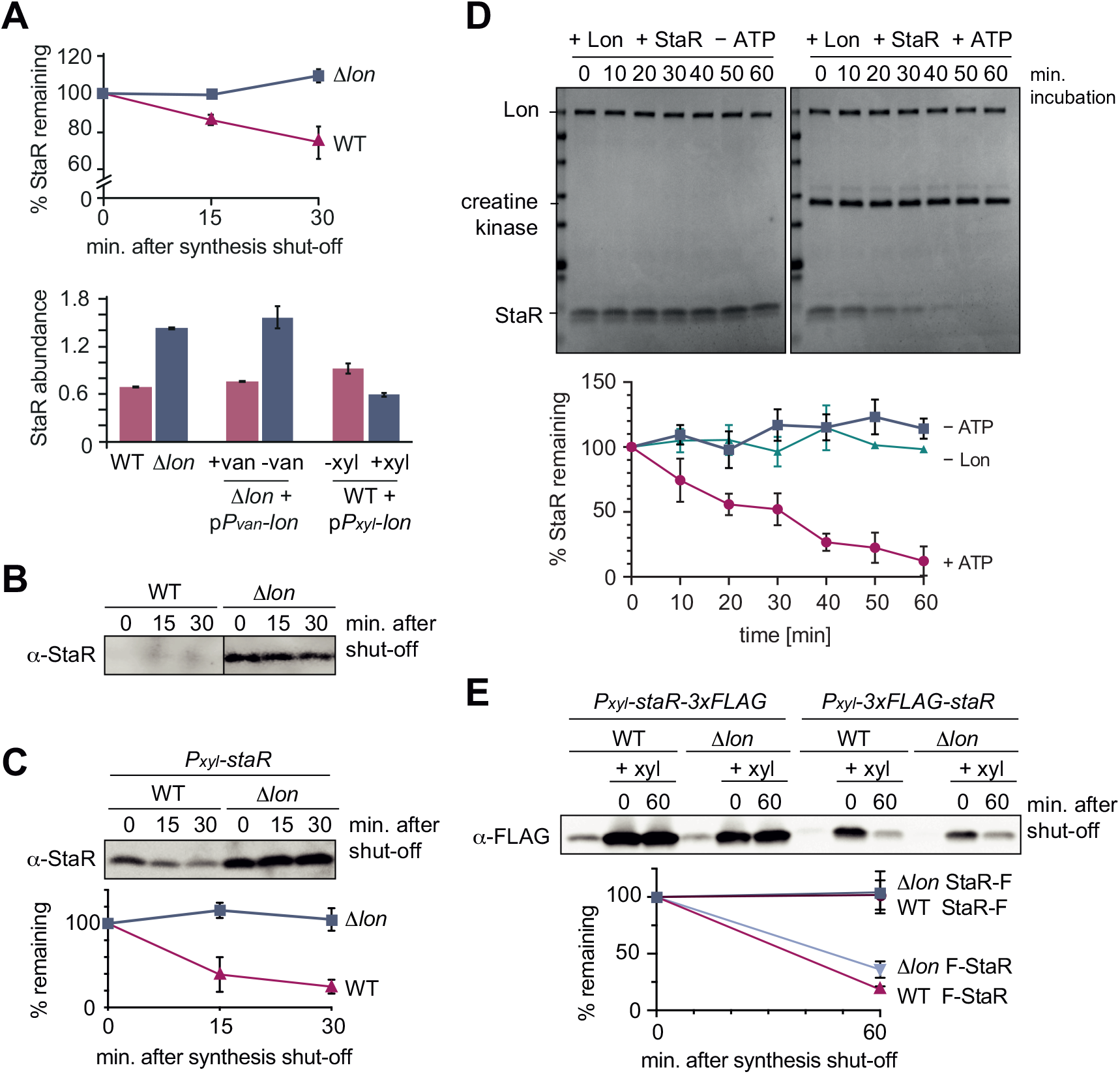
The developmental regulator StaR is a Lon substrate. (A) Proteomics data obtained for StaR. The upper graph shows StaR stability in WT and *Δlon* (LS2382) cells and the lower graph StaR abundance in the different strain backgrounds and conditions as determined by mass spectrometry. Each data point represents the mean protein abundance of the two experimental replicates, error bars show standard deviations. (B) *In vivo* degradation assay of StaR at native expression levels in WT and *Δlon* (KJ546) cells. (C) *In vivo* degradation assay of StaR after xylose-induced *staR* overexpression in WT and *Δlon* (KJ546) backgrounds. The graph shows mean values and standard deviations of relative protein levels after protein synthesis shut-off determined from three independent experiments. (D) *In vitro* assay showing degradation of StaR by Lon. 4 µM StaR and 0.125 µM Lon hexamer was incubated in the presence (+ATP) or absence (−ATP, control) of an ATP regeneration system. The graph shows the relative StaR levels normalized to Lon or CK (−Lon sample) levels of three independent experiments and are represented as means with standard deviations. (E) N-terminally and C-terminally tagged StaR (3xFLAG-StaR and StaR-3xFLAG, respectively) were expressed ectopically by xylose induction and their protein levels were assessed in wild type and *Δlon* (KJ546) cells prior to induction (first lanes of each set of three) and after induction either before (t = 0 min) or after shutting-off protein synthesis (t = 60 min). The graph shows the mean values of relative protein levels and standard deviations of three independent experiments.

We also investigated if Lon recognizes StaR via a degron sequence at one of the termini and monitored the degradation of FLAG-tagged StaR variants. Addition of the 3xFLAG tag to the C-terminus of StaR (StaR-F) completely abolished degradation (Fig. 2E), indicating that a freely accessible C-terminus of StaR is required for degradation by Lon. Addition of the tag to the N-terminus of StaR (F-StaR) still enabled notable degradation within 60 min after shutting down protein synthesis (Fig. 2E). However, degradation of this N-terminally tagged StaR was also observed in the *Δlon* strain, suggesting that another protease degrades this StaR variant, probably because of changes in StaR folding that result from the addition of the tag. These data show that native N-and C-termini of the protein are required for Lon-dependent degradation.

### Lon ensures cell cycle-dependent regulation of StaR abundance

As a transcriptional regulator of stalk biogenesis and holdfast production, StaR function is expected to be particularly needed at the beginning of the cell cycle, when the swarmer cell differentiates into a stalked cell (Fig. 3A). Indeed, previously published RNA sequencing data show that *staR* mRNA levels fluctuate during the cell cycle, peaking in the swarmer and early stalked cell before declining during S-phase and remaining low until cell division (Fig. 3B) [30, 31]. Furthermore, existing ribosome profiling data show that StaR is translated predominantly in the swarmer and early stalked cells, but only at low levels during the remaining cell cycle [30, 31]. Quantification of StaR protein levels by Western blotting in synchronized cultures showed that protein levels follow this expression pattern in wild type cultures (Fig. 3C). StaR was detectable within 30 min after synchronization before it was strongly downregulated and remained below the limit of detection for the rest of the cell cycle. Strikingly, the cell cycle-dependent changes in protein abundance were absent in the *Δlon* strain, in which StaR levels remained high until 75 minutes after synchronization (Fig. 3C). Thus, Lon-dependent degradation of StaR is required for establishing oscillations of StaR levels during the cell cycle.

**Figure 3.**
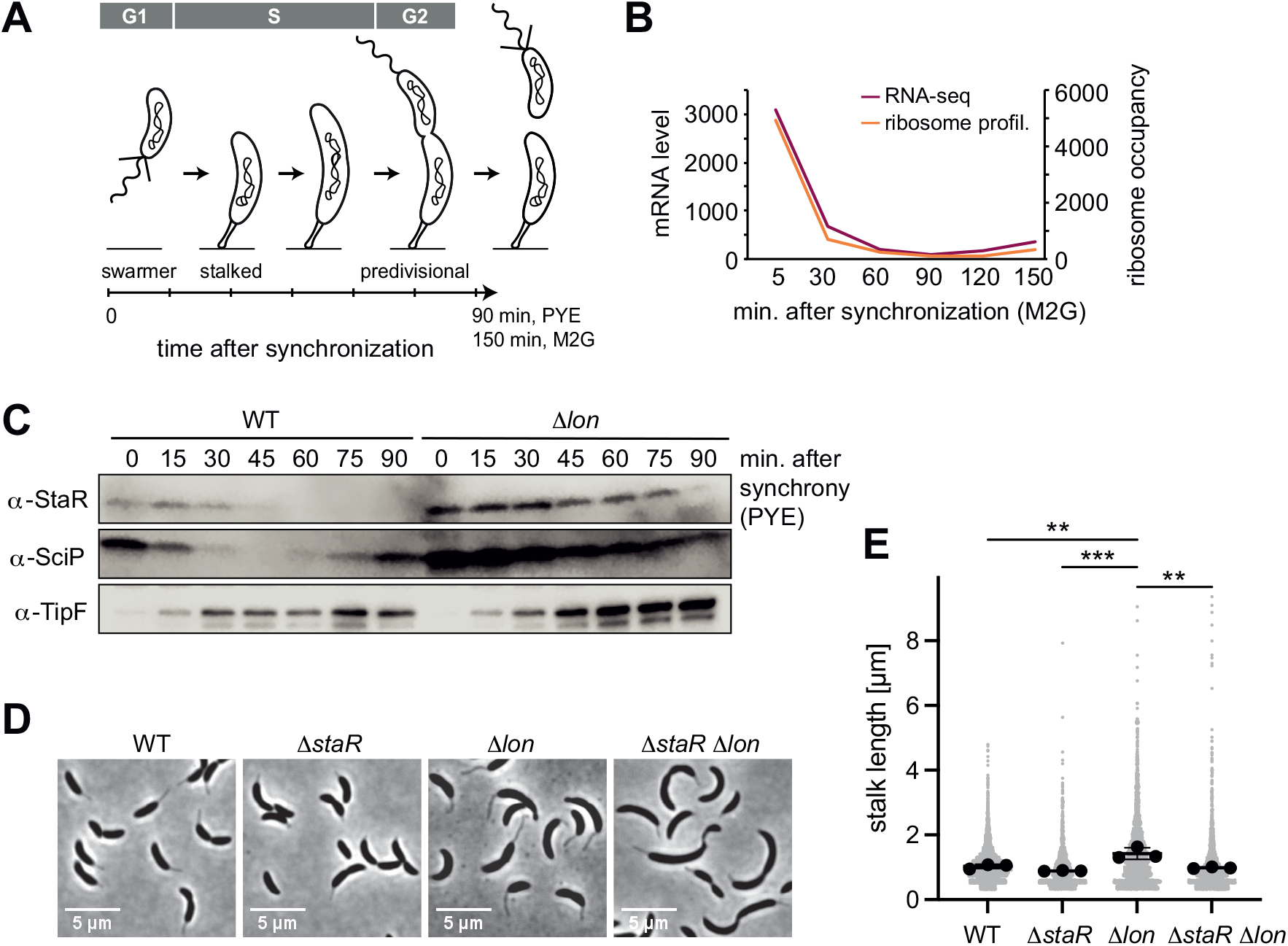
Lon ensures cell cycle-dependent accumulation of StaR and proper stalk length regulation. (A) Schematic illustration of the *C. crescentus* cell cycle and how the different cell cycle phases are associated with distinct morphological states. The time line indicates how the cell cycle progresses over time. (B) RNA-sequencing and ribosome profiling data for *staR*, as published previously [30, 31]. (C) Protein levels of StaR, SciP and TipF in synchronized wild type and *Δlon* (KJ546) cultures over 90 min following release of swarmer cells into PYE medium. Western blots for SciP and TipF were included as controls for proteins with well-characterized cell cycle patterns [62, 63]. (D) Phase contrast microscopy images depicting morphological differences between *C. crescentus* wild type (WT) the single mutants *ΔstaR* and *Δlon* (KJ546) and the double mutant *ΔstaRΔlon* when grown in PYE at 30 °C. (E) Quantifications of stalk length under optimal conditions (PYE, 30°C) of the strains shown in (D). *N* was at least 1 800 total for each strain obtained from three biological replicates. Statistical significance was determined by ordinary one-way ANOVA (Tukeýs multiple comparisons test: WT vs *Δlon p* = 0.0047, **; *ΔstaR* vs *Δlon p* = 0.0007,***; *Δlon* vs *ΔstaRΔlon p* =0.0025, **).

### Lon-mediated StaR proteolysis is required for proper stalk biogenesis

Next, we assessed the importance of Lon-mediated StaR proteolysis for correct stalk biogenesis. Consistent with a previous study [23], we observed that the stalks of *Δlon* cells are significantly elongated compared to the wild type (Fig. 3D-E). Because StaR regulates stalk length, holdfast production and other developmental processes [32, 33], we reasoned that the abnormal stalk length of *Δlon* cells might be caused by the higher abundance of StaR in these cells. To address this hypothesis, we introduced the *ΔstaR* deletion into the *Δlon* strain background and assessed stalk length of this *ΔstaRΔlon* double mutant (Fig. 3D-E, Fig. S1). The *ΔstaRΔlon* mutant phenocopied the *ΔstaR* single mutant, in which stalks are shortened compared to *Δlon* cells, demonstrating that Lon affects stalk length through StaR (Fig. 3D-E). To further investigate the relationship between Lon, StaR and stalk length we also analyzed stalk morphology under phosphate limiting conditions, in which stalks are drastically elongated in *C. crescentus* [34]. Stalk length in the different strain backgrounds followed the same trend as under optimal conditions (Fig. S1). However, all four strains were able to strongly elongate their stalks under phosphate starvation, suggesting that StaR and Lon are not required for starvation-dependent stalk elongation. Taken together, our data demonstrate that Lon-mediated degradation of StaR is required for proper stalk biogenesis during the cell cycle.

### The two putative Lon substrates CCNA_00944 and MotD are part of one single protein that corresponds to FliK

In addition to StaR, our proteomics experiments identified several proteins involved in flagella-based motility and chemotaxis as potential Lon substrates (Fig. 1F, Dataset S1). Particularly promising hits in this group of proteins were CCNA_00944 and MotD (CCNA_00945) that are encoded by partly overlapping open reading frames and showed similar changes in protein abundance and stability in the Lon deficient and Lon overproducing strains in our proteomics experiments (Fig. 4A-B). While CCNA_00944 is annotated as a flagella hook length determination protein, MotD is annotated as a chemotaxis protein. According to a signature based annotation of MotD in UniProtKB (entry A0A0H3C6R3), MotD contains a conserved Flg hook domain that is commonly present in the C-terminal portion of FliK proteins that control flagella hook length in many bacteria [35]. When we attempted to clone CCNA_00944, we repeatedly observed one additional guanosine in the cloned gene sequence that was not present in the reference genome sequence of *C. crescentus* NA1000, the strain that we use. This insertion, which is also present in the sequence reads of previously published RNA-sequencing data [36], generates a frameshift that merges the CCNA_00944 gene with the downstream located *motD* gene, thus forming one single open reading frame, of which the 3’ portion corresponds to *motD* (Fig. 4B). This resulting gene corresponds to a single open reading frame (CC_0900) in *C. crescentus* CB15, the isolate from which NA1000 is derived. These observations indicate that CCNA_00944 and *motD* are incorrectly annotated in the reference genome of NA1000 and instead form one continuous open reading frame. Consistently, when we performed Western blot analysis with antiserum raised against the protein portion corresponding to MotD we detected one single protein band that runs at high molecular weight (Fig. S2), confirming that CCNA_00944 and *motD* form together one single open reading frame. Because the C-terminal portion of this new gene encodes a Flg hook domain that is a characteristic of FliK proteins in other bacteria and because no other gene has so far been annotated as *fliK* in *C. crescentus*, we named the new gene *fliK* (Fig. 4B). This annotation of *fliK* is also in line with a recent study in *Sinorhizobium meliloti*, which suggested that the *motD* gene from alpha-proteobacteria should be renamed *fliK* [37].

**Figure 4.**
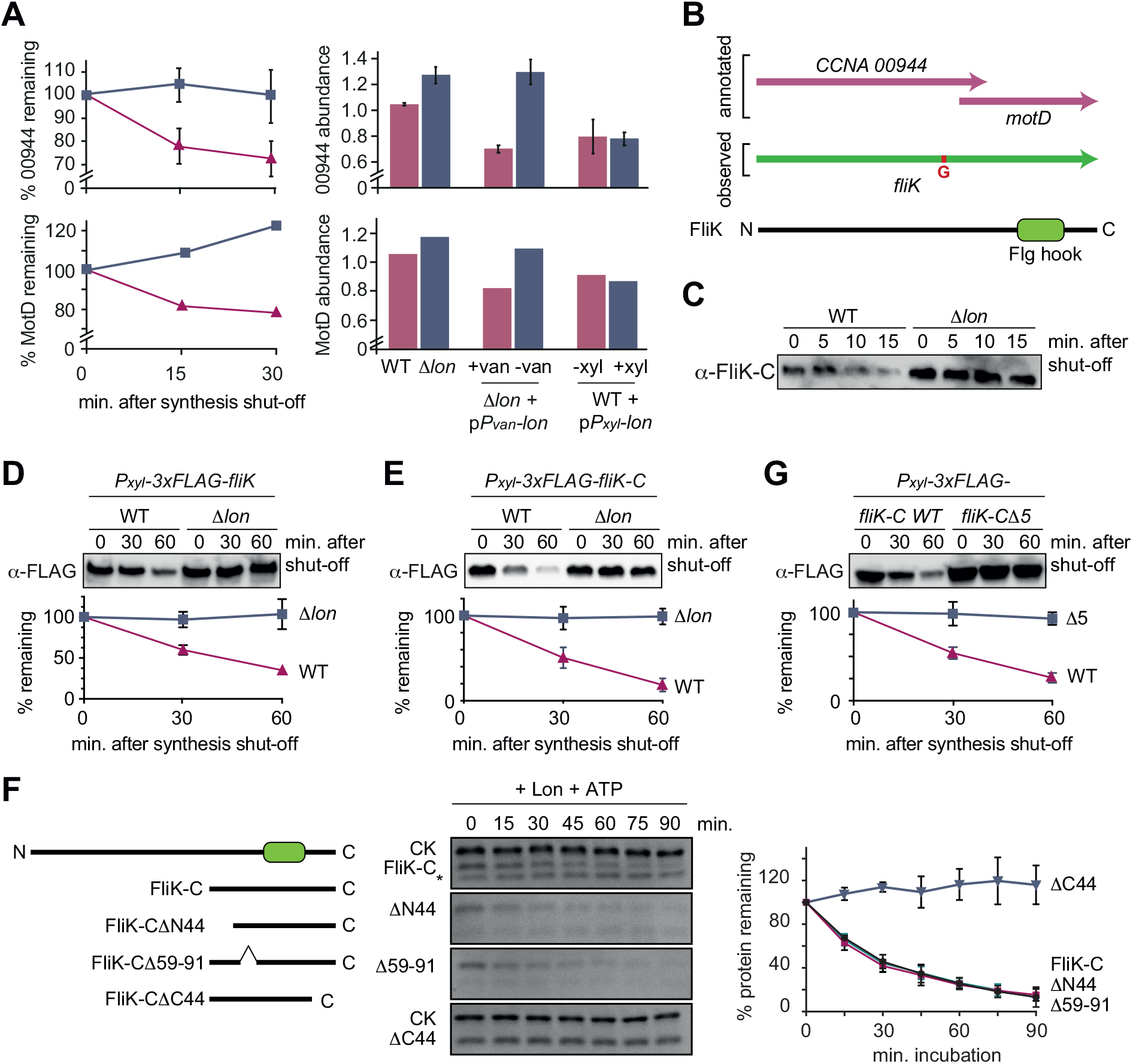
The flagella hook length regulator FliK is a Lon substrate with a C-terminal degradation tag. (A) Proteomics data obtained for CCNA_00944 and CCNA_00945 (MotD). The graphs on the left show CCNA_00944 and MotD stability in WT and *Δlon* (LS2382) cells, and graphs on the right show CCNA_00944 and MotD abundance in the different strain backgrounds and conditions as determined by mass spectrometry. Each data point for CCNA_00944 represents the mean protein abundance of the two experimental replicates, error bars show standard deviations. MotD was only detected in one of the replicates. (B) Schematic representation of the CCNA_00944 and *motD* (CCNA_00945) genes as annotated in the *C. crescentus* NA1000 genome, and how the presence of an additional guanosine (highlighted in red) merges the two genes to form one continues gene that was named *fliK*. The corresponding FliK protein contains a Flg hook domain. (C) *In vivo* degradation assay of full-length FliK in wild type (WT) and *Δlon* (KJ546) cells. Samples were taken 0, 5, 10 and 15 min after shutting off protein synthesis. (D) *In vivo* degradation assay of ectopically expressed N-terminally 3xFLAG-tagged full-length FliK in wild type (WT) and *Δlon* (KJ546) cells. The graph shows mean values and standard deviation of relative protein levels 0, 30 and 60 min after protein synthesis shut-off determined from three independent experiments. (E) *In vivo* degradation assay of the C-terminal part of FliK (FliK-C) in wild type (WT) and *Δlon* (KJ546) cells. N-terminally 3xFLAG tagged FliK-C was ectopically expressed and samples were taken at indicated time points after shut-off of protein synthesis. Quantifications show mean values obtained from three independent biological replicates and error bars represent standard deviation. (F) *In vitro* degradation assays showing Lon-dependent degradation of FliK-C and truncated FliK-C proteins lacking N-terminal, internal or C-terminal regions, graphically illustrated on the left side of the panel. Degradation assays were carried out in Lon reaction buffer with 4 µM of one of the FliK-C variants, 0.125 µM of Lon hexamer in the presence of the ATP regeneration system (ATP, creatine phosphate, creatine kinase [CK]). The band intensities of the FliK-C variants from three independent experiments are represented as means with standard deviations on the right side of the panel. (G) *In vivo* degradation assay of N-terminally 3xFLAG tagged FliK-C (WT) and a variant lacking the C-terminal five amino acids (FliK-CΔ5) after xylose induction in wild type cells. Samples were taken at indicated time points after shut-off of protein synthesis. Quantifications show mean values of relative protein levels obtained from three biological replicates and error bars represent standard deviation.

### FliK is a Lon substrate that is recognized at its C-terminus

Having established that CCNA_00944 and MotD are part of the same FliK protein, we next investigated its regulation by Lon. Consistent with our proteomics data, we found that FliK abundance and stability were notably increased in *Δlon* cells, consistent with FliK being a Lon substrate (Fig. 4C). To investigate the sequence determinants required for Lon to interact with FliK, we expressed an N-terminally FLAG tagged version of FliK and monitored its stability *in vivo*. Similar to the non-tagged FliK, 3xFLAG-FliK was efficiently degraded in wild type cells but stable in cells lacking Lon (Fig. 4D). This result led us to hypothesize that the recognition by Lon occurs via the C-terminal domain of FliK. Therefore, we monitored the stability of the C-terminal portion of FliK containing the Flg hook domain, which corresponds to the formerly annotated MotD protein. Like the full-length protein, degradation of this truncated FliK protein (FliK-C), with a FLAG-tag at the N-terminus, depended strongly on Lon (Fig. 4E). Furthermore, *in vitro* degradation assays showed that Lon degrades non-tagged FliK-C in an ATP-dependent manner (Fig. 4F, Fig. S3). Based on these results, we conclude that FliK is a Lon substrate, and that its C-terminal portion containing the Flg hook domain is sufficient for Lon-dependent degradation.

To further pinpoint the regions within FliK-C that are required for Lon-dependent turnover, we analyzed the degradation of a set of additional truncation mutants. According to the MobiDB database [38], the C-terminal portion of FliK lists two unordered regions. We engineered FliK-C variants that lack either of these unordered regions (FliK-CΔN44 and FliK-CΔ59-91) or the C-terminal part (FliK-CΔC44). *In vitro* degradation assays showed that deletion of the unordered regions did not influence degradation, whereas removal of the 44 C-terminal amino acids completely stabilized the protein (Fig. 4F). This result, and the fact that some of the degrons recognized by Lon are located at the very C-terminus of Lon substrates [14, 39–41], prompted us to determine the *in vivo* stability of a FliK-C variant lacking only the C-terminal five amino acids with the sequence LDIRI (3xFLAG-FliK-CΔ5) (Fig. 4G). This deletion abolished FliK-C degradation (Fig. 4G), demonstrating that the interaction between Lon and FliK depends on these C-terminal amino acids.

### Lon ensures temporally restricted accumulation of FliK during the cell cycle

Like many other proteins involved in flagella biogenesis in *C. crescentus*, the transcription of the two annotated genes CCNA_00944 and *motD* that together form the *fliK* gene is cell cycle regulated [30, 31], with mRNA levels and ribosome occupancy peaking in late S-phase when a new flagellum at the pole opposite the stalk is being assembled (Fig. 5A). Consistently, Western blot analysis with synchronized wild type *C. crescentus* cultures showed that FliK protein was not detectable in the beginning of the cell cycle, but began to accumulate in late stalked cells before reaching a maximum in abundance in predivisional cells shortly before cell division (Fig. 5B). This cell cycle-dependent pattern of FliK abundance was completely absent in the *Δlon* strain, in which FliK was already detectable in swarmer cells and remained at high levels throughout the cell cycle (Fig. 5B). This result shows that, as in the case of StaR, Lon is absolutely necessary to ensure that the protein is eliminated in the cell cycle phase when its function is no longer needed.

**Figure 5.**
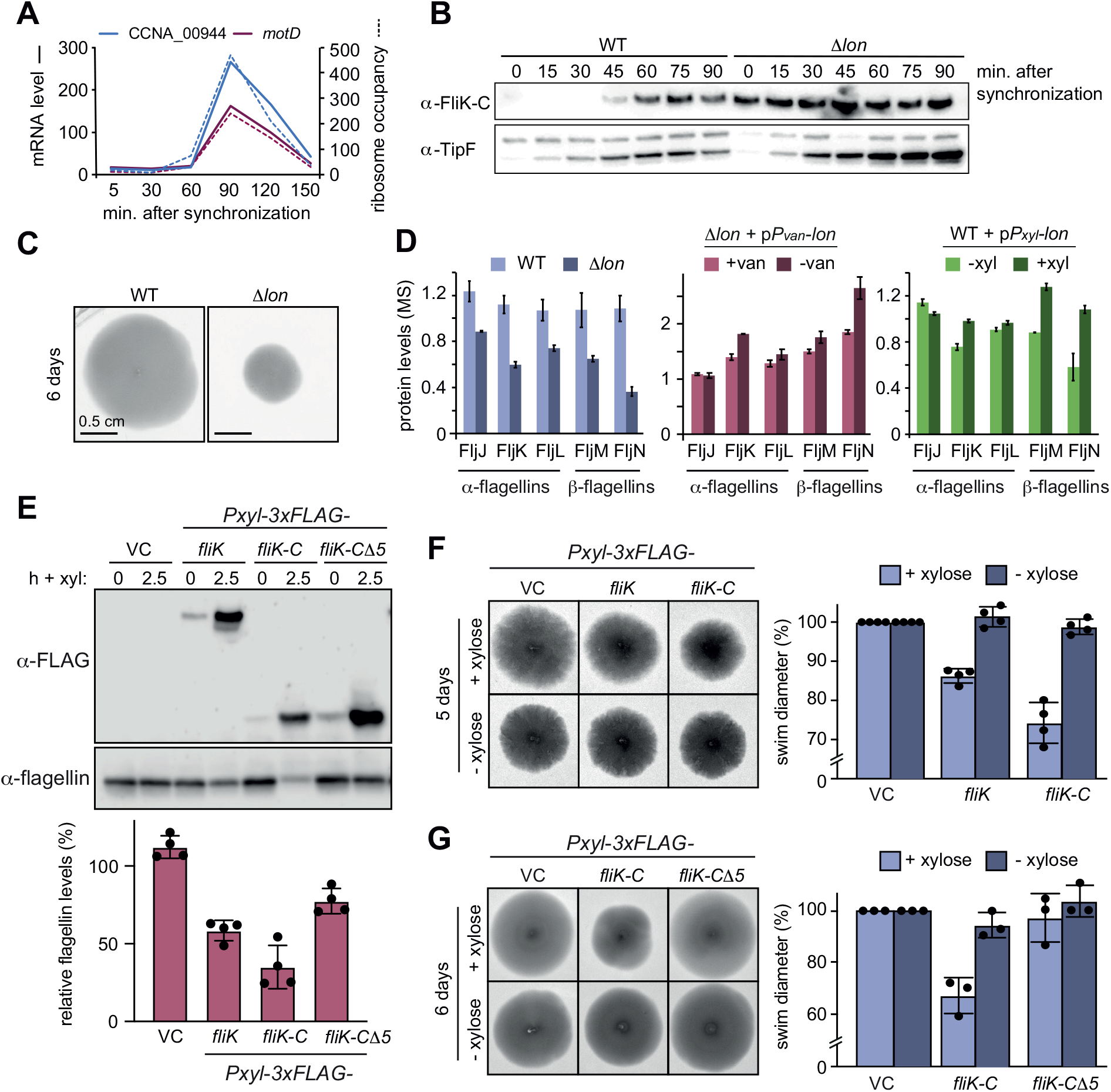
Lon-dependent degradation ensures temporal regulation of FliK levels during the cell cycle, which is needed for normal flagellin expression and motility. (A) RNA-sequencing and ribosome profiling data for CCNA_00944 and *motD* (CCNA_00945), as previously published [30, 31]. (B) Protein levels of FliK and TipF in synchronized wild type and *Δlon* (KJ546) cultures over 90 min following release of swarmer cells into PYE medium. TipF was included as a control. (C) Motility assay of *C. crescentus* wild type (WT) and the *Δlon* (KJ546) mutant in PYE soft agar after 6 days. (D) Flagellin protein levels as determined by mass spectrometry in wild type and *Δlon* (LS2382) mutant cells as well as in the other strain backgrounds and conditions, see Fig. 1. (E) Western blots showing flagellin levels of strains harboring the empty vector (VC) or plasmids for expression of 3xFLAG-tagged FliK, FliK-C or FliK-CΔ5 before (0) and after induction of expression by xylose for 2.5 hours. Induction of the respective FliK variants was determined using an anti-FLAG antibody. The graph shows the relative flagellin levels compared to the uninduced condition (without xylose) for each strain, as determined by four independent experiments, mean values and standard deviations are indicated. (F) Motility assay in soft agar of strains overexpressing 3xFLAG-tagged FliK and FliK-C by xylose induction (+xylose) in comparison to the vector control (VC) and non-inducing conditions (−xylose). The graph shows the relative swim diameters from four biological replicates (VC was set to 100 %), means and standard deviations are indicated. (G) Motility assay in soft agar of strains overexpressing 3xFLAG-tagged FliK-C and FliK-CΔ5 by xylose induction (+xylose) in comparison to the vector control (VC) and non-inducing conditions (−xylose). The graph shows the relative swim diameters from three biological replicates (VC was set to 100 %), means and standard deviations are indicated.

### Precise regulation of FliK abundance is required for proper flagellin expression and flagella function

Based on our results that Lon degrades proteins involved in flagella biosynthesis and the observation that *Δlon* cells have a severe motility defect in soft agar compared to the wild type (Fig. 5C) [27], we hypothesized that Lon-mediated degradation impacts *C. crescentus* motility. In line with this idea, our proteomics data revealed that the *α*-flagellins FljJ, FljK and FljL and the *β*-flagellins FljM and FljN, which compose the structural components of flagella, are misregulated in the *Δlon* mutant, as well as in the Lon depletion and overexpression strains (Fig. 5D, Dataset S1). Cytoplasmic flagellin levels were previously shown to be affected by FliK homologs in other bacteria [37]. To test if the higher abundance of FliK in *Δlon* cells may contribute to the observed misregulation of flagellin expression, we overexpressed FLAG-tagged FliK from a medium copy vector and found that *α*-flagellins levels were downregulated to approximately 55 % compared to the uninduced control (Fig. 5E). Interestingly, overexpression of the FliK-C variant caused an even stronger reduction in *α*-flagellin levels to 30 %. This effect was dependent on the last C-terminal five amino acids, since overexpression of the FliK-C*Δ*5 variant affected *α*-flagellin levels only mildly (Fig. 5E). These results demonstrate that the C-terminal part of FliK mediates the effect on flagellin levels, probably through an interaction with the secretion gate FlhB that is known to depend on the five C-terminal amino acids of FliK [42, 43].

To test if the FliK-mediated effect on flagellin expression translates to changes in motility, we measured the swim diameter of cells overexpressing FLAG-tagged FliK, FliK-C and FliK-C*Δ*5 (Fig. 5F-G). Consistent with the observed lower flagellin levels, overexpression of FliK led to a reproducible reduction in swim diameter compared to the uninduced control condition, which again was clearly exacerbated in the strain overexpressing FliK-C (Fig. 5F). Conversely, FliK-C*Δ*5 overexpression had no effect on motility (Fig. 5G). Together, these data demonstrate that oversupply of FliK results in reduced flagellin levels as well as compromised motility, and that these effects are mediated by the same C-terminal amino acids of FliK that are also required for its degradation. Thus, the reduced flagellin levels and motility defects of *Δlon* can at least in part be attributed to the stabilization of FliK in these cells.

## Discussion

In all cells the concentrations of specific proteins must be precisely regulated to maintain cellular functions and to orchestrate complex cellular behaviors in response to external and internal cues. This study uncovered a novel role of the highly conserved protease Lon in coordinating cell differentiation with cell cycle processes in the dimorphic bacterium *C. crescentus*. In this bacterium each cell cycle phase is coupled to a distinct morphological state [16]. This coupling of cell differentiation with core cell cycle events requires sophisticated mechanisms that coordinate these processes in space and time. Previous work established that in *C. crescentus* large sets of genes are transcriptionally regulated in a cell cycle-dependent manner [17, 30, 44]. Using a proteomics approach, we found that several proteins encoded by cell cycle-regulated genes are Lon substrates. The identified Lon substrates include important regulators and structural components required for *C. crescentus* development and cell cycle progression. Our results show that active proteolysis of at least some of these proteins by Lon is required to rapidly clear these proteins following a cell cycle-dependent decrease in their transcription, thus restricting their accumulation to the cell cycle phase when their function is needed (Fig. 6). Although the catalytic activity of Lon does not seem to change during the cell cycle under optimal conditions [40], it is possible that the degradation of these proteins is affected by their functional state, for example by their binding to DNA or interactions with other proteins. Indeed, previous work showed that degradation of both SciP and CcrM is modulated by DNA binding [22, 40], and in the case of the AAA+ ATPase DnaA, ATP binding seems to increase protein stability [45, 46].

**Figure 6.**
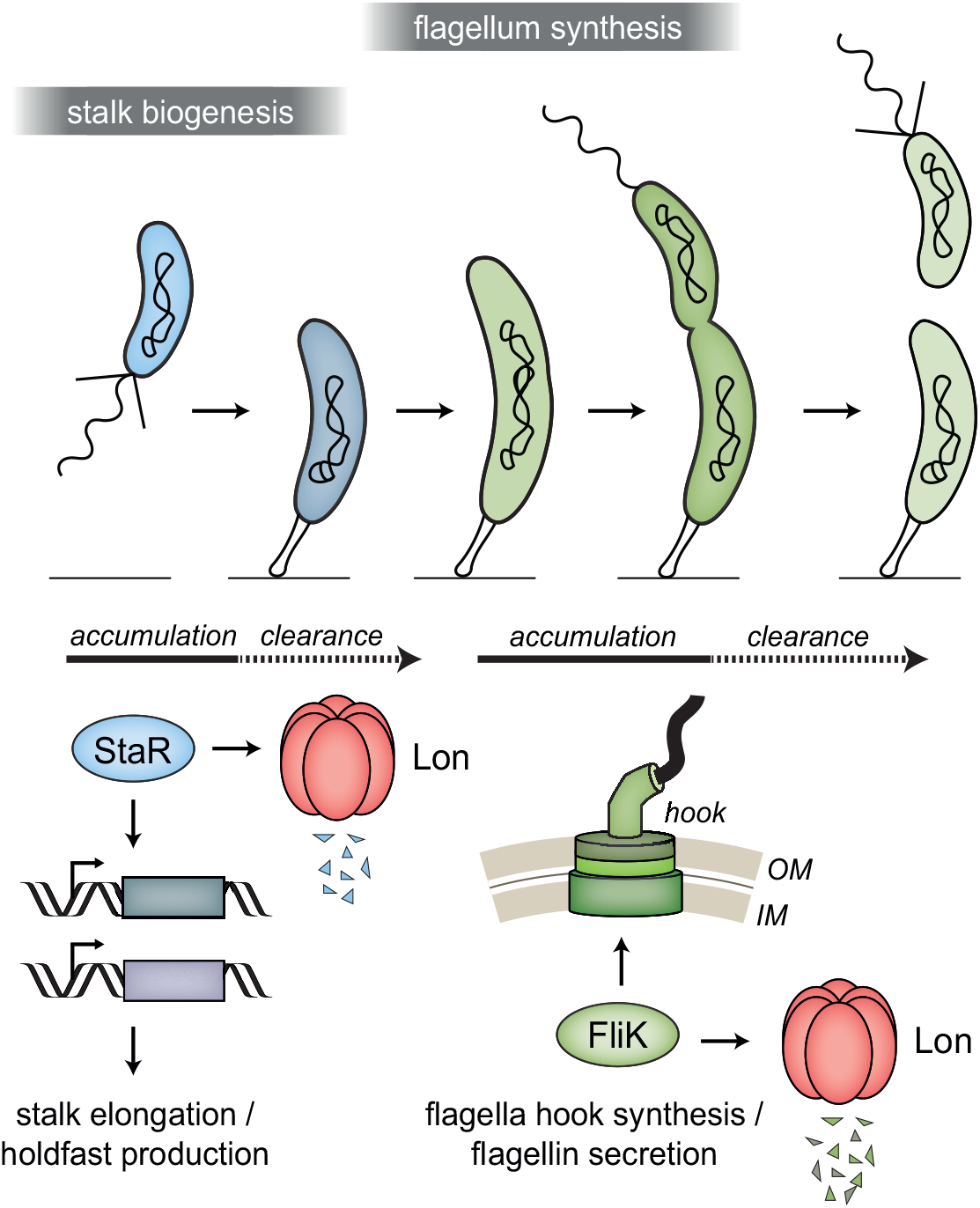
Lon ensures temporal regulation of stalk and flagella biogenesis during the *C. crescentus* cell cycle. Lon specifically degrades StaR, a transcriptional regulator of stalk biogenesis and holdfast production, and FliK, a protein involved in regulating flagella hook synthesis. The expression of *staR* peaks in swarmer cells, while the expression of *fliK* peaks in late stalked and predivisional cells [17, 30, 31]. Our study shows that Lon-dependent proteolysis is required to rapidly eliminate these proteins when their expression levels drop, thus outpacing synthesis. This ensures that StaR abundance is temporally restricted to the swarmer-to-stalked cell transition (shown in blue) and FliK abundance to the late stalked and predivisional cell (shown in green) when their functions are needed for stalk biogenesis or flagella synthesis, respectively.

We focused our studies on the stalk regulator StaR and the flagella regulator FliK, which were among the proteins whose abundance and stability was most strongly affected by Lon. Like DnaA, CcrM and SciP, the transcriptional regulator StaR is a DNA-binding protein. It was initially identified as a regulator of stalk biogenesis [32] and was later shown to regulate holdfast development by directly inhibiting the expression of the holdfast inhibitor HfiA [33]; holdfast is a polysaccharide-rich adhesin that is produced at the nascent stalked cell pole in late swarmer cells and allows *C. crescentus* to attach to surfaces [16]. Stalk biogenesis and surface attachment must be tightly regulated during the cell cycle, in particular under environmental conditions [33], and our work revealed that Lon-mediated proteolysis contributes to this by regulating the stability of StaR and likely other proteins involved in this process, such as the histidine phosphotransferase ShpA and the polysaccharide biosynthesis protein CCNA_02361 that we identified as putative Lon substrates in our proteomics screen (Fig. 1F). The gene encoding CCNA_02361 (CC_2278), which was previously shown to contribute to surface attachment [47], shows a similar cell cycle-dependent pattern in mRNA levels and ribosome occupancy as *staR* [30, 31], with the highest mRNA abundance and translation rate in the swarmer state (Fig. 1F). Thus, Lon may also contribute to temporally regulating the abundance of this protein during the cell cycle.

In addition to proteins involved in stalk biogenesis and surface attachment, we identified several proteins required for flagella-mediated motility and chemotaxis as novel Lon substrates (Fig. 1F), and investigated FliK in detail. FliK proteins are thought to function as molecular rulers that precisely monitor flagellar hook assembly and mediate a switch from export of hook protein to filament protein, the final stage of flagella synthesis, once the hook has reached a certain length [48, 49]. Although previous work made significant progress in understanding the molecular function of this important protein in enterobacteria, proteolytic control mechanisms contributing to FliK regulation have so far not been described. Moreover, in bacteria outside the gamma-proteobacteria FliK remains poorly studied. In *C. crescentus* a *fliK* gene has previously not been annotated, and our new data revealed that *fliK* corresponds to a gene that was previously annotated as two separate but overlapping genes. We demonstrate that Lon-mediated proteolysis ensures cell cycle-dependent accumulation of FliK during late S-phase, when the cell prepares for cell division by building a new flagellum and chemotaxis apparatus at the swarmer pole (Fig. 6), and that this precise regulation of FliK abundance is critical for correct flagellin levels and normal motility. Our data uncovered a critical role of the C-terminal amino acids of FliK for both proteolytic control and FliK function, the latter likely depending on an interaction with the secretion protein FlhB [42, 43, 50], indicating a tight coupling between FliK function and degradation.

Interestingly, in addition to the flagella hook length regulator FliK, our data indicate that the flagella hook protein FlgE itself is regulated by Lon in *C. crescentus*. FlgE was among the proteins that satisfied two criteria in our proteomics approach, and we verified by Western blotting that FlgE degradation partly depends on Lon (Dataset S1, Fig. S4). This finding is also consistent with previous work showing that Lon degrades FlgE in *E. coli* [12], and reinforces the previously made notion that precise regulation of the cellular concentrations of FlgE as well as FliK is an important requirement for the correct temporal order of flagella assembly [51]. In several other bacteria, including important human pathogens, Lon has been linked to flagella-mediated motility [52–55]. In many of these cases the nature of substrates mediating the observed Lon-dependent effects on motility remains unknown. However, in *Bacillus subtilis* it was shown that Lon specifically degrades the master regulator of flagellar biosynthesis SwrA by a mechanism requiring the adaptor SmiA [56]. In contrast to SwrA, both StaR and FliK are robustly degraded by Lon *in vitro* (Fig. 2D, Fig. 4F, Fig. S3). While these data suggest that no adaptors or other accessory proteins are required to mediate the interaction between these proteins and Lon, it is possible that additional factors exist that modulate the rate of Lon-mediated proteolysis of these proteins in response to specific conditions.

In conclusion, our work provides new insights into the cellular roles of Lon and emphasizes the importance of proteolysis in adjusting the amounts of regulatory proteins involved in critical cellular processes, including cell differentiation and cell cycle progression. Importantly, in addition to StaR and FliK our work identified many other proteins as putative Lon substrates and the precise role of Lon in the regulation of these proteins will be worthwhile to investigate in detail in future studies. Furthermore, our work highlights quantitative proteomics using isobaric mass tags as a powerful approach for the identification of novel protease substrates that could be exploited to identify candidate substrate under diverse growth conditions, in different species or of other proteases.

## Materials and Methods

### Strains and plasmids

All bacterial strains, plasmids and primers used in this study are listed in Table S1.

### Plasmid construction

#### Expression plasmids for protein purification

Plasmids used for protein expression are based on the pSUMO-YHRC backbone and were constructed as follows: The coding sequences of the *staR* (pMF56-c88) and *fliK*-C (formerly *motD*; pMF61) genes were amplified from *C. crescentus* NA1000 genomic DNA with the primers listed in Table S1 (see sheets listing primers and vector fragments for sequences and primer combinations, respectively). The backbone vector pSUMO-YHRC was amplified in two parts disrupting the kanamycin resistance gene in order to reduce background (using primer pairs oMJF34/oMJF36 and oMJF37/oMJF38, Table S1). Following the PCR, the template was digested with DpnI (10 U) and the remaining PCR fragments subsequently purified by gel extraction. Fragments were then assembled using Gibson assembly [57]. Vectors containing deletions of an annotated gene (FliK-C truncations: pMF66, pMF67-A, pMF68-A) were derived from vectors harboring the full-length coding sequence in a similar manner using the primer pairs specified in Table S1.

#### Replicating plasmids

pDJO145 (pBX-MCS-4 containing *NdeI-3xFLAG-KpnI*): Plasmid pBX-MCS-4 [58] was amplified using primers oDJO13 and oDJO41. The sequence encoding the triple FLAG tag (*3xFLAG*; GAC TAC AAA GAC CAT GAC GGT GAT TAT AAA GAT CAT GAC ATC GAC TAC AAG GAC GAC GAC GAC AAG) was amplified from a plasmid using primers oDJO42 and oDJO43 adding a KpnI-site followed by a stop codon to the 3’end of *3xFLAG*. The two amplified fragments were then joined by Gibson assembly [57].

pDJO151 (pBX-MCS-4 containing P*xyl*-*staR-3xFLAG*):

*staR* was amplified with primers oDJO44 and oDJO45 using chromosomal *C. crescentus* NA1000 DNA as template and cloned into NdeI-cut pDJO145 using Gibson assembly.

pDJO157 (pBX-MCS-4 containing P*xyl-3xFLAG-staR*):

*staR* was amplified with primers oDJO46 and oDJO47 using chromosomal *C. crescentus* NA1000 DNA as template and cloned into KpnI-cut pDJO145 using Gibson assembly.

pDJO173 (pBX-MCS-4 containing *Pxyl-flgE-3xFLAG*):

*flgE* was amplified with primers oDJO65 and oDJO66 using chromosomal *C. crescentus* NA1000 DNA as template and cloned into NdeI-cut pDJO145 using Gibson assembly.

pDJO200 (pBX-MCS-4 containing P*xyl-3xFLAG-fliK-C*):

*flik-C (CCNA_00945)* was amplified with primers oDJO87 and oDJO88 using chromosomal *C. crescentus* NA1000 DNA as template and cloned into KpnI-cut pDJO145 using Gibson assembly.

pDJO410 (pBX-MCS-4 containing P*xyl*-3xFLAG*-fliK-C*D*5*):

*flik-CD5* was amplified with primers oDJO87 and oDJO179 using *C. crescentus* NA1000 DNA as template and cloned into KpnI-cut pDJO145 using Gibson assembly.

pDJO487 (pBX-MCS-4 containing P*xyl-3xFLAG-fliK)*:

*fliK* was amplified with primers oDJO75 and oDJO88 using chromosomal *C. crescentus* NA1000 DNA as template and cloned into KpnI-cut pDJO145 using Gibson assembly.

### Strain construction

To generate the *ΔstaRΔlon* strain (KJ1037), the *Δ*s*taR* deletion was introduced into the *Δlon* strain (KJ546) by two-step recombination [59] after transformation with plasmid pNTPS138-*ΔstaR* (pAF491, [33]). Briefly, transformants were selected on kanamycin plates, single colonies were grown overnight in PYE and plated on PYE containing sucrose. Single sucrose-resistant colonies were subsequently screened for kanamycin sensitivity and the *staR* knockout was confirmed by colony PCR using primers oDJO40 and oDJO38.

*C. crescentus* strains carrying replicating plasmids were created by transforming the plasmids into the respective strain backgrounds by electroporation.

### Standard growth conditions

*C. crescentus* strains were routinely grown at 30°C in PYE medium while shaking at 200 rpm and, if necessary, regularly diluted to assure growth in the exponential phase. If required, the medium was supplemented with xylose (0.3 % final), glucose (0.2 % final) or vanillate (500 µM final). Antibiotics were used at following concentration (liquid/solid media): gentamycin 0.625/5 µg/ml, chloramphenicol 1/1 µg/ml, oxytetracycline 1/2 µg/ml. Experiments were generally performed in the absence of antibiotic when using strains in which the resistance cassette was integrated into the chromosome. For phosphate starvation experiments, log-phase cells grown in M2G (minimal medium with 0.2 % glucose) were washed and transferred to M5G medium lacking phosphate.

*E. coli* strains for cloning purposes were grown in LB medium at 37 °C, supplemented with antibiotics at following concentrations (liquid/solid media): chloramphenicol 20/40 µg/ml, gentamycin 15/20 µg/ml, kanamycin 30/50 µg/ml, oxytetracyclin 12/12 µg/ml.

### Synchronization of *C. crescentus* cultures

To synchronize *C. crescentus* cultures, cells were pelleted by centrifugation at 8000 rpm for 4 minutes at 4 °C. The supernatant was aspirated, and tubes kept on ice. Pellets were resuspended in 1 ml of 1x M2 salts on ice. 1 ml of cold Percoll was added as samples were mixed well. The mixture was aliquoted into two Eppendorf tubes and centrifuged at 10000 ×g for 20 minutes at 4 °C. The top layer of the cells was aspirated, and the swarmer cells were moved into a new tube. Swarmer cells were washed twice with 1 ml of cold 1x M2 salts and finally resuspended in 20 ml prewarmed PYE medium containing antibiotic if required. Samples were taken immediately after resuspending the cell pellet and subsequently at the indicated time points for immunoblot analysis.

### Immunoblot analysis

For whole cell extract analysis, 1 ml culture samples were collected after the indicated treatments and time points, and cell pellets were obtained by centrifugation. Cell pellets were resuspended in appropriate amounts of 1x SDS sample buffer, to ensure normalization of the samples by units OD_600_ of the cultures. Samples were boiled at 98 °C for 10 min and frozen at −20 °C until further use. The thawed samples were separated by SDS-PAGE, and subsequently transferred to nitrocellulose membranes by a semi-dry blotting procedure as per manufacturer guidelines. The stain-free Tris-glycine gels (Bio-Rad) used enabled the assessment of equal loading of total protein in-gel as well as the quality of the transfers.

Membranes were blocked in 10 % skim milk powder in TBS-Tween (TBST) and protein levels were detected using the following primary antibodies and dilutions in 3 % skim milk powder in TBST: anti-CcrM 1:5 000 [60]; anti-DnaA 1:5 000 [61], anti-Lon 1:10 000 (kindly provided by R.T. Sauer), anti-FLAG M2 antibody 1:5 000 (Sigma), anti-SciP 1:2 000 [62], anti-flagellin 1:2 000 (kindly provided by Y. Brun), anti-StaR 1:500 [33], anti-TipF 1:5 000 (kindly provided by P. Viollier), [63]. Secondary antibodies, 1:5 000 dilutions of anti-rabbit or anti-mouse HRP-conjugated antibodies (Thermo Scientific), and SuperSignal Femto West (Thermo Fisher Scientific) were used to detect primary antibody binding. Immunoblots were scanned using a Chemidoc (Bio-Rad) system or a LI-COR Odyssey Fc system. Relative signal intensities were quantified using the Image Lab software package (Bio-Rad) or ImageJ software.

### *In vivo* degradation assay

To assess protein degradation *in vivo*, cells were grown under the appropriate conditions (e.g., for 1 to 2 hours in the presence of xylose to induce expression of FLAG-tagged proteins), and subsequently protein synthesis was shut-off by addition of chloramphenicol (100 μg/ml). Samples were taken at the indicated time points and snap frozen in liquid nitrogen before preparation for analysis by Western blotting.

### Quantitative proteomics

Sample preparation was performed as previously described [64]. In brief, two independent cultures for each analyzed condition were harvested by centrifugation. Cell pellets were washed using cold ddH2O and stored at −80 °C. Protein digestion, TMT10 plex isobaric labeling and the mass spectrometrical analysis were performed by the Clinical Proteomics Mass Spectrometry facility, Karolinska Institute/Karolinska University Hospital/Science for Life Laboratory.

To identify putative Lon substrates, first, the protein abundances for each condition within one biological replicate were considered to calculate the following ratios (see Dataset S1): protein levels in *Δlon* at t = 0 min/protein levels in WT at t = 0 min (*Δlon* 0/WT 0) to identify proteins with a higher steady state level in the absence of Lon; protein levels after Lon depletion/protein levels before Lon depletion (Lon depletion (no van)/*lon* expression (+ van)) to identify differences in protein levels after Lon depletion; protein levels before *lon* overexpression in the presence of glucose/protein levels after 1 h xylose-induced *lon* overexpression (gluc/+ xyl 1h *lon* overexpression), to identify proteins that are down-regulated by *lon* overexpression; protein levels in WT at t = 0 min/ protein levels in WT at t = 30 min (WT 0/WT 30), to identify proteins that are degraded in wild type cells after protein synthesis shut-off. Additionally, we calculated the ratios of protein levels in *Δlon* cells at t = 0 min/protein levels in *Δlon* cells at t = 30 min after protein synthesis shut-off (*Δlon* 0/*Δlon* 30). Subsequently, the average of the ratios obtained from the two biological replicates was calculated and used for further analysis. Some proteins, as indicated in Dataset S1, were detected in only one of the replicate data sets. In these cases, only the ratio from the replicate, in which they were detected, was considered.

Next, we selected all proteins with a ratio of (*Δlon* 0/WT 0) ≥ 1.1 for the group “higher levels in *Δlon*”. Similarly, we selected all proteins with a ratio of (Lon depletion (no van)/ *lon* expression (+ van)) ≥ 1.1 for the group “higher levels after Lon depletion”. For the group “lower levels after *lon* overexpression” we selected all proteins with a ratio of (gluc/+ xyl 1h *lon* overexpression) ≥ 1.05. For the group “stabilized in *Δlon*” we first selected all proteins with a ratio of (WT 0/WT 30) ≥ 1. For those, we calculated the ratio of (WT 0/ WT 30)/(*Δlon* 0/*Δlon* 30) and chose the proteins with a ratio ≥ 1.05. Finally, we used jvenn [65] to determine the overlap between the groups (Fig. 1D). Proteins that fulfilled 4 or 3 categories were assigned to functional categories as previously described [64].

### Protein purification

Protein purification was adapted from Holmberg e*t al*. 2014 [66]. In brief, BL21-SI/pCodonPlus cells were transformed using a pSUMO-YHRC derived vector by electroporation. Transformants were selected on LB agar plates lacking NaCl (LBON) supplemented with kanamycin and chloramphenicol and pre-cultures inoculated with 20 colonies before being cultivated at 30 °C overnight. Pre-cultures were diluted 1:100 in 1 l 2xYTON and grown to OD 1.0–1.5. Protein expression was induced with 0.5 mM IPTG and 0.2 M NaCl either for 4.5 h at either 30°C (StaR and FliK-C) or overnight at 20 °C (in case of FliK-C truncations). After centrifugation, cell pellets were stored dry at −80 °C.

Pellets were resuspended in HK500MG (40 mM HEPES-KOH pH7.5, 500 mM KCl, 5 mM MgCl, 5% Glycerol) supplemented with 1 mM PMSF, 1 mg/ml Lysozyme and 3 µl Benzonase/10 ml suspension and topped up to 20 ml. Cells were then lysed by 2–4 passes through an EmulsiFlex-C3 high-pressure homogenizer. Lysate was cleared by centrifugation at 32 500 ×g at 4 °C for 0.5–1 h. The protein of interest was bound to 1 g Protino Ni-IDA beads at 4°C/on ice for 30 minutes. After washing 5 times with approx. 45 ml HK500MG protein was eluted using HK500MG + 250 mM Imidazole and fractions with protein concentrations ≥1 mg/ml were pooled. For 6xHis-SUMO tag removal, 4 µg/ml Ulp1-6xHis was added and imidazole was removed in parallel by dialysis against HK500MG. Tag depletion (except for StaR) was achieved by binding to 1 g Protino Ni-IDA beads as before and flow through containing purified protein was collected. Protein concentration was then checked via SDS-PAGE (Bio-Rad 4–20 % Mini-PROTEAN® TGX Stain-Free™ protein gel) and InstantBlue protein stain (Expedeon) and quantified using Bio-Rad ImageLab 6.0.1. Protein samples were aliquoted and stored at −80 °C.

### StaR refolding

Because StaR was not soluble in neither of the tested buffer conditions and formed precipitates, the protein was refolded (adapted from [67, 68]). The precipitate was collected (centrifugation at 7 197 ×g, 4°C) and solubilized in buffer S (50 mM HEPES pH8.0, 6 M guanidinium-HCl, 1 mM EDTA, 10 mM DTT) and the protein concentration was adjusted to 0.2 mg/ml. The protein solution was then diluted with an equal volume dialysis buffer D1 (50 mM HEPES pH8.0, 2 M guanidinium-HCl, 2 mM EDTA) and dialyzed against 125 volumes dialysis buffer D1 followed by dialysis against 125 volumes dialysis buffer D2 (50 mM HEPES pH8.0, 1 M guanidinium-HCl, 2 mM EDTA, 0,4 M Sucrose, 500 mM KCl, 2 mM DTT). The dialysis buffer was then diluted with one buffer volume of dialysis buffer D3 (50 mM HEPES pH8.0, 2 mM EDTA, 0,4 M Sucrose, 500 mM KCl, 2 mM DTT) and dialyzed. This was followed by a final dialysis against 125 volumes of dialysis buffer D3 to remove remaining guanidinium-HCl. Each dialysis step was carried out at 4 °C for approx. 24 h.

Afterwards, insoluble StaR molecules were removed by centrifugation (20 000 ×g, 4°C, 10 min) and cleared refolded StaR was supplemented with additional 2 mM DTT and concentrated using a centrifugal filter with a MWCO of 3 kDa (VWR #516-0227P) and stored at −80 °C. The final concentration was determined by SDS-PAGE using a BSA standard and visualized by InstantBlue protein stain (Expedeon).

### FliK-C antibody production

Purified FliK-C was used as antigen to generate rabbit polyclonal antisera (Davids Biotechnologie GmbH).

### *In vitro* degradation assays

*In vitro* degradation assays were performed as published previously [24]. The reaction was carried out in Lon reaction buffer (25 mM Tris pH8.0, 100 mM KCl, 10 mM MgCl_2_, 1 mM DTT) employing 0.75 µM Lon (0.125 µM Lon_6_), 4 µM substrate (if not stated otherwise) and an ATP regeneration system (4 mM ATP, 15 mM creatine phosphate, 75 µg/ml creatine kinase). The reaction and the ATP regeneration system were prepared separately pre-warmed to 30 °C (approx. 4 min). The reaction was started by adding the ATP regeneration system. Samples were taken at indicated time points and quenched by 1 vol. 2x SDS sample buffer (120 mM Tris-Cl pH 6.8, 4 % SDS, 20 % glycerol, 0.02 % bromophenol blue) and snap frozen in liquid nitrogen. Samples were boiled at 65 °C for 10 minutes and separated by SDS-Page (Bio-Rad 4–20 % Mini-PROTEAN® TGX Stain-Free™ protein gel), visualized by InstantBlue protein stain (Expedeon) and quantified using Bio-Rad ImageLab 6.0.1. Substrate levels were normalized to Lon and/or creatine kinase levels (in case of –Lon samples).

### Microscopy

Cells were fixed by addition of formaldehyde (1 % final) to culture samples and stored at 4 °C. For visualization, fixed cells were transferred onto 1 % agarose pads attached to glass slides, covered with a coverslip and transferred to the microscope. A T*i* eclipse inverted research microscope (Nikon) with 100x/1.45 numerical aperture (NA) objective (Nikon) was used to collect phase-contrast images. The images were processed with Fiji (ImageJ).

### Stalk length measurements

To quantify the stalk length of cells, microscopy images were analyzed using ImageJ. Briefly, the scale was set to 1 pixel representing an equivalent of 0.0646 μm. The stalk was then manually selected and length was determined (Fig. S1). For half-automated analysis (Fig. 3E), the software BacStalk was used [69]. The data files containing microscopic images were added to the program, the channel “phase contrast” was selected, and the scale set to 1 px ≙ 0.0646 μm. Stalk detection was activated by the setting “My cells have stalks”. For cell detection the default settings were used.

### Motility assays

To assess motility, strains were grown in PYE media, supplemented with gentamycin to maintain replicating plasmids when necessary, and cultures were diluted to an OD_600_ of 0.1. Subsequently, 1 µl of each sample was injected about 2 mm vertically into PYE agar plates (0.35 %), supplemented with gentamycin or gentamycin and xylose when indicated, using a pipette. The plates were incubated at 30 °C and pictures were taken every day with the setting “Blots: Colorimetric” using a ChemiDoc (BioRAD).

## Acknowledgements

We thank Peter Chien and his lab for sharing aliquots of purified Lon and for their help with Lon purifications, members of the Jonas lab for discussions and specifically Roya Akar for technical assistance, Claes Andréasson and his lab for providing the BL21-SI/pCodonPlus strain and the pSUMO-YHRC vector and for their help with the His-SUMO protein purification procedures, Yves Brun and Patrick Viollier for sharing aliquots of antibodies and Sean Crosson for providing plasmids and the *ΔstaR* strain. We also thank the Clinical proteomics facility at KI/KS for support and advice as well as the Protein Expression and Characterization facility at SciLifeLab for sharing their equipment and providing help with the protein purifications. The study was financially supported by grants from the Swedish Research Council (Dnr. 2016-03300 and 2020-03545), a future leaders grant from the Swedish Foundation for Strategic Research (FFL15-0005) and funding from the Strategic Research Area (SFO) program distributed through Stockholm University.

## Supplemental Information

**Figure S1.**
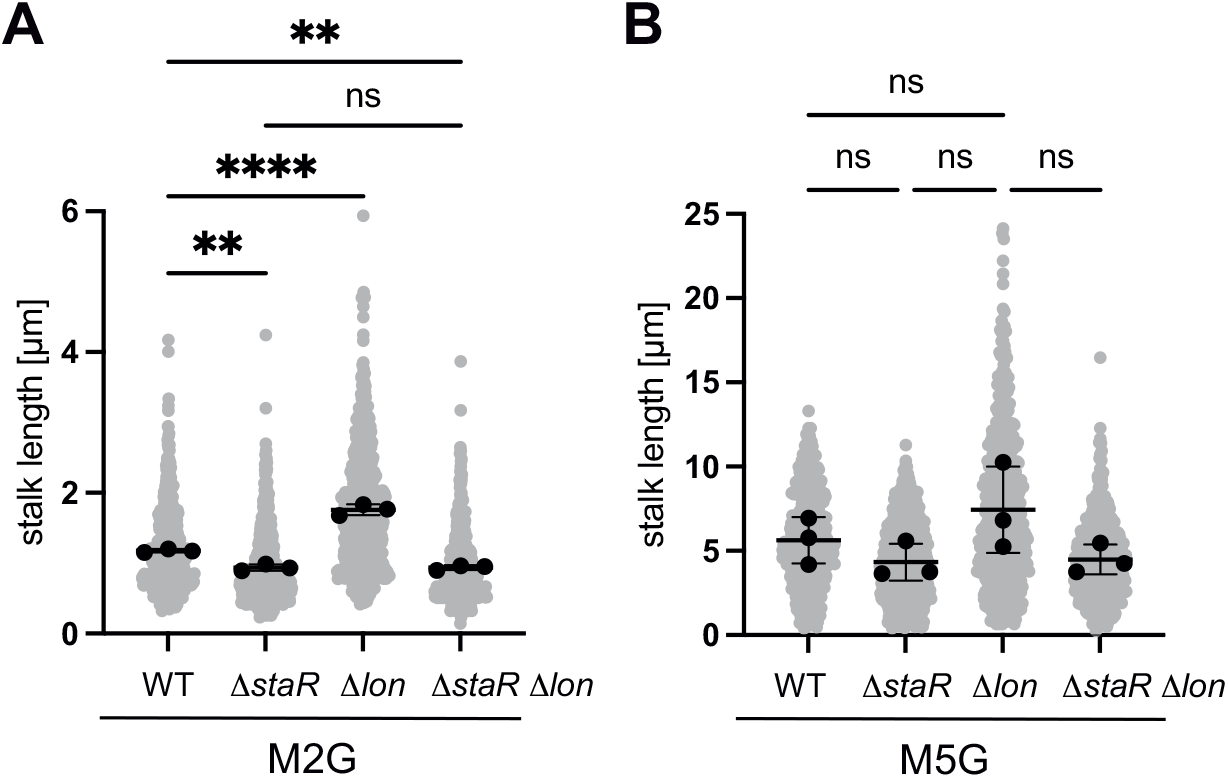
Quantification of stalk length under standard (M2G) and phosphate-limiting conditions (M5G). (A) Quantification of stalk length of the indicated strains grown in M2G medium. *N* was at least 400 total for each strain and obtained from three biological replicates. Statistical significance was determined by ordinary one-way ANOVA (Tukeýs multiple comparisons test: WT vs Δ*staR p* = 0.0018, *\*\*;* WT vs Δ*lon* (KJ546) *p* < 0.0001*, ****;* WT vs Δ*staR*Δ*lon p* = 0.0018*, **;* Δ*staR* vs Δ*staR*Δ*lon,* ns (not significant). (*B*) Cells of the indicated strain backgrounds were pre-grown in M2G medium, washed, and subsequently grown for 24 hours in the phosphate-limiting medium M5G. *N* was at least 400 total for each strain and obtained from three biological replicates. The differences between the strains were statistically not significant as determined by ordinary one-way ANOVA (Tukeýs multiple comparisons test).

**Figure S2.**
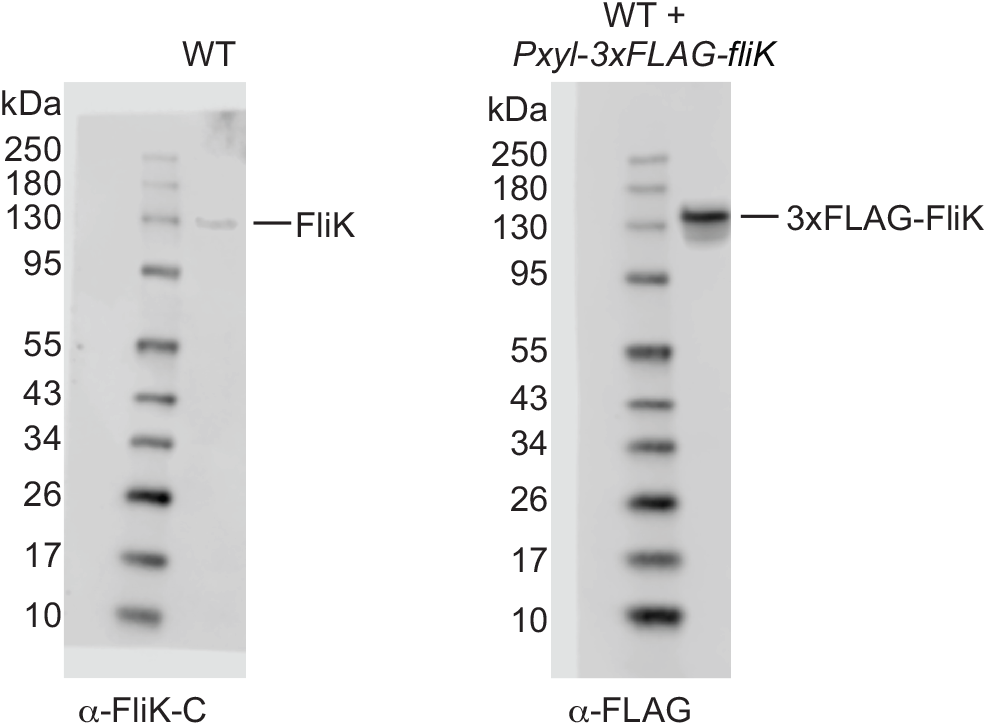
Western blots of native FliK and 3xFLAG tagged FliK. Left panel: native FliK was detected using anti-FliK-C antibodies in wild type (WT) cell extracts. The overlay of the blot with the picture of the prestained protein standard is shown, indicating that native FliK migrates with the 130 kDa standard band. Right panel: 3xFLAG tagged FliK was induced by xylose addition in wild type (WT) cells and detected using anti-FLAG antibodies in extracts as indicated. The overlay of the blot with the picture of the prestained protein standard is shown, indicating that 3xFLAG tagged FliK migrates slightly above the 130 kDa standard band.

**Figure S3.**
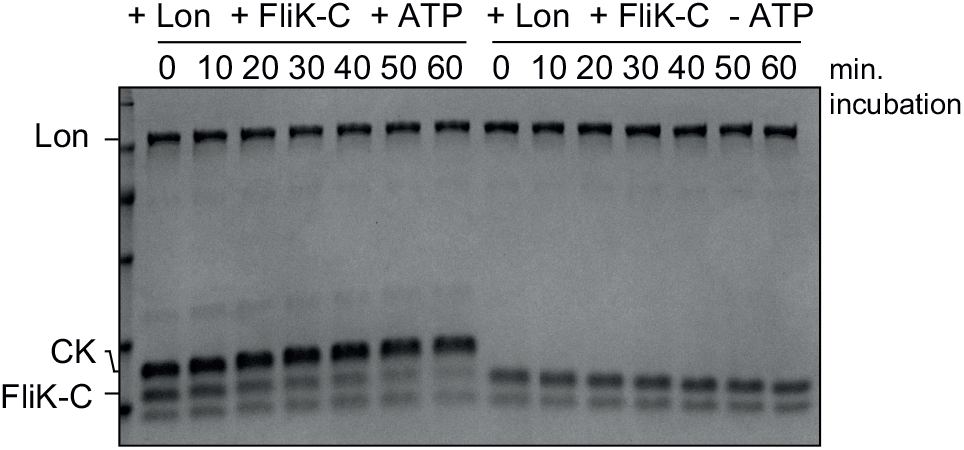
*In vitro* FliK-C degradation by Lon is ATP dependent. The *in vitro* degradation assay was carried out containing 4 µM FliK-C and 0.125 µm Lon hexamer in the presence (+ATP) or absence (−ATP) of the ATP regeneration system (ATP, creatine phosphate and creatine kinase [CK]). A representative gel of two independent replicates is shown.

**Figure S4:**
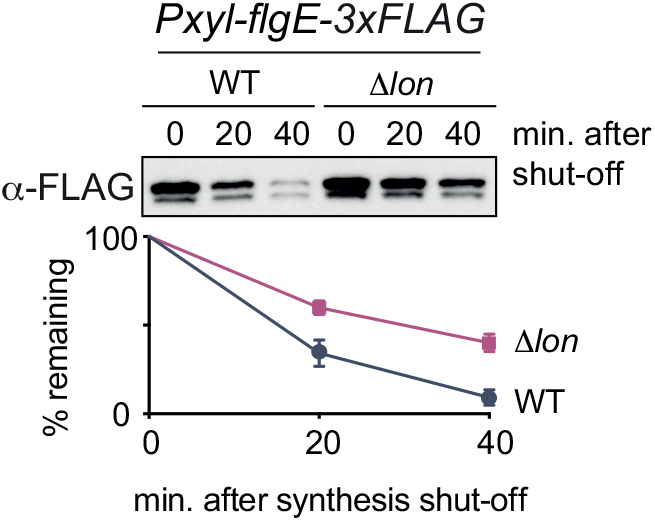
FlgE degradation is partially dependent on Lon. *In vivo* degradation assay of C-terminally FLAG tagged FlgE (FlgE-3×FLAG) after overexpression by xylose induction in wild type (WT) and Δ*lon* (KJ546) cells. Samples were taken at the indicated time points after protein synthesis shut-off. Mean values and standard deviation of relative protein levels after protein synthesis shut-off were determined from three independent experiments.

**Dataset S1. Proteomics data.**

**Table S1. Bacterial strains, plasmids and primers used in this study.**

